# Generation of human bronchial organoids for SARS-CoV-2 research

**DOI:** 10.1101/2020.05.25.115600

**Authors:** Tatsuya Suzuki, Yumi Itoh, Yusuke Sakai, Akatsuki Saito, Daisuke Okuzaki, Daisuke Motooka, Shohei Minami, Takeshi Kobayashi, Takuya Yamamoto, Toru Okamoto, Kazuo Takayama

## Abstract

Coronavirus disease 2019 (COVID-19) is a disease that causes fatal disorders including severe pneumonia. To develop a therapeutic drug for COVID-19, a model that can reproduce the viral life cycle and evaluate the drug efficacy of anti-viral drugs is essential. In this study, we established a method to generate human bronchial organoids (hBO) from commercially available cryopreserved human bronchial epithelial cells and examined whether they could be used as a model for severe acute respiratory syndrome coronavirus 2 (SARS-CoV-2) research. Our hBO contain basal, club, ciliated, and goblet cells. Angiotensin-converting enzyme 2 (ACE2), which is a receptor for SARS-CoV-2, and transmembrane serine proteinase 2 (TMPRSS2), which is an essential serine protease for priming spike (S) protein of SARS-CoV-2, were highly expressed. After SARS-CoV-2 infection, not only the intracellular viral genome, but also progeny virus, cytotoxicity, pyknotic cells, and moderate increases of the type I interferon signal could be observed. Treatment with camostat, an inhibitor of TMPRSS2, reduced the viral copy number to 2% of the control group. Furthermore, the gene expression profile in SARS-CoV-2-infected hBO was obtained by performing RNA-seq analysis. In conclusion, we succeeded in generating hBO that can be used for SARS-CoV-2 research and COVID-19 drug discovery.

**Graphical abstract:** 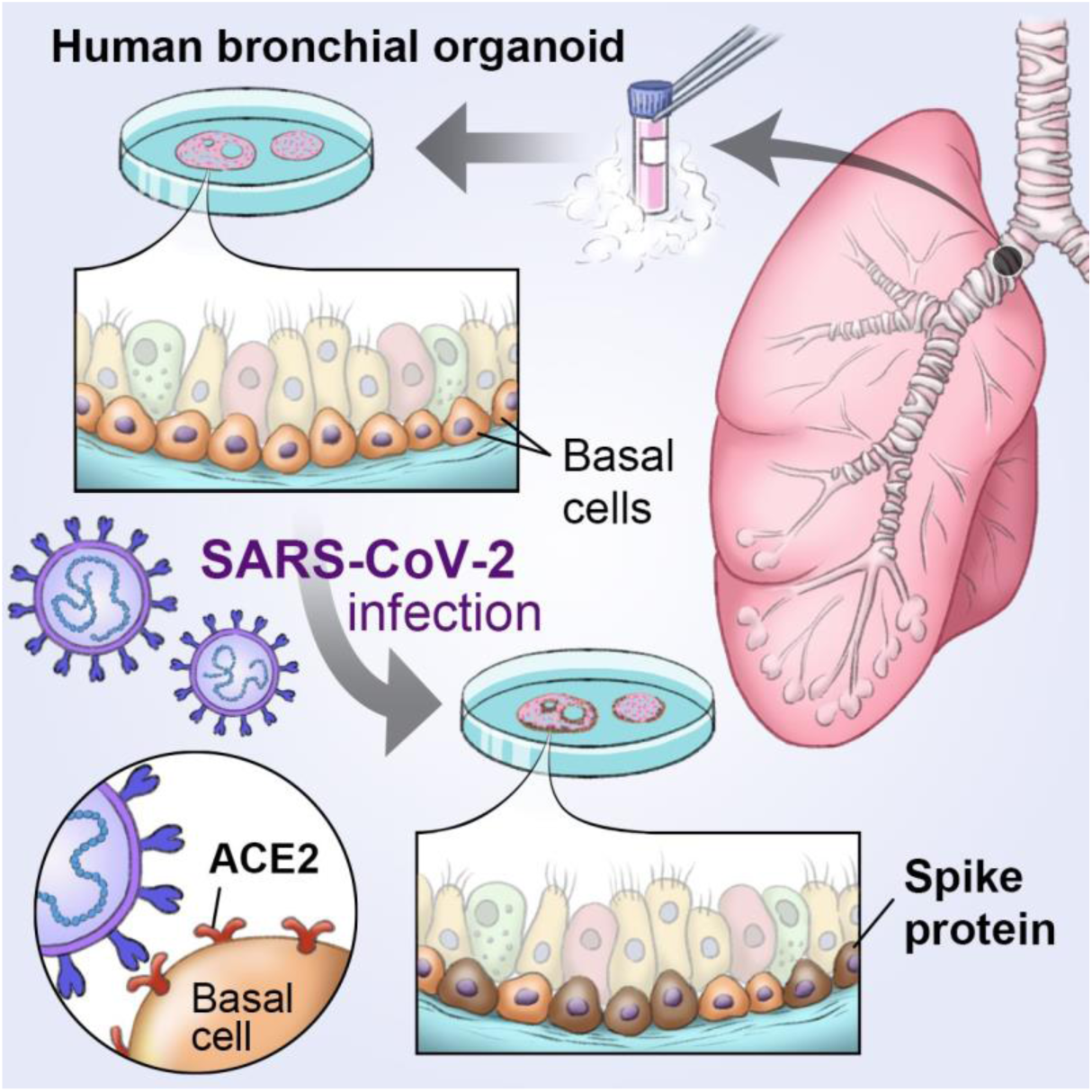

## Introduction

The “2019-new coronavirus disease (COVID-19) was first reported in China in December 2019 ^1^ and declared a pandemic by the WHO in March 2020 ^2^. Severe pneumonia is most frequently observed in COVID-19 patients, and the number of COVID-19 patients and deaths are still increasing. These conditions have made it difficult for research on severe acute respiratory syndrome coronavirus 2 (SARS-CoV-2), which is the causative virus of COVID-19, to keep pace. SARS-CoV-2 is composed of four proteins: S (spike), E (envelope), M (membrane), and N (nucleocapsid) proteins. It is known that angiotensin-converting enzyme 2 (ACE2) is a SARS-CoV-2 receptor, and transmembrane serine proteinase 2 (TMPRSS2) is essential for priming S protein ^3^. Thus, to accelerate SARS-CoV-2 research, a novel lung model that reproduces the viral life cycle with intact expression of these host factors is indispensable.

A number of animal and cell models that can be used for SARS-CoV-2 research have been reported (Takayama et al., in revision), but an *in vitro* lung model that can evaluate candidate therapeutic agents for COVID-19 is essential for conducting large-scale drug screening. Human lung organoids are excellent tools that can faithfully mimic the lung functions of living organisms ^4^. The lungs consist of bronchi and alveoli. Lukassen et al. have performed single-cell analysis of alveoli and bronchi and reported that ACE2 is predominantly expressed in the transient secretory cell type, which displays a transient cell state between goblet and ciliated cells, in bronchi, but TMPRSS2 is strongly expressed in both lung tissues ^5^. Therefore, a bronchial organoid containing transient secretory, goblet, and ciliated cells could be a useful model for SARS-CoV-2 research.

Several reports have verified the usefulness of two-dimensional (2D) culturing airway epithelial cells in SARS-CoV2 studies. For example, SARS-CoV-2 infection experiments using human bronchial epithelial cells (hBEpC) showed cytopathic effects 96 hr after the infection on the layers of human airway epithelial cells ^6^. In addition, hBEpC cultured using an air-liquid interface culture system can be used to evaluate viral infection, replication, and the drug efficacy of remdesivir ^7^. 2D culture systems of hBEpC are relatively easy to use, but they cannot reproduce the cellular microenvironment in the living body and are difficult to use for long-time culture. Recently, it was shown that SARS-CoV-2 can infect and replicate in human pluripotent stem cell (PSC)-derived lung organoids containing bronchial epithelial cells and alveolar epithelial cells ^8^. However, these organoids exhibit a fetal phenotype rather than an adult type ^9,10^. Adult-type bronchial organoids are essential because of the severe infection caused by COVID-19 in adults. Human bronchial organoids (hBO) with adult phenotype can be established from intact human lung tissue ^11^. However, it is difficult for many researchers to obtain an intact lung biopsy sample, because the process requires the approval of an ethics committee and informed consent from the donor. Therefore, in this study, we developed a method for generating hBO from commercially available cryopreserved adult hBEpC and applied it to SARS-CoV-2 research.

## Results

### Generation of human bronchial organoids from cryopreserved adult bronchial epithelial cells

We searched for the conditions that could establish hBO from cryopreserved adult hBEpC. We found that after embedding the hBEpC in Matrigel and culturing with advanced DMEM/F12 medium containing FGF2, FGF7, FGF10, Noggin, R-spondin 1, Y-27632, and SB202190 (expansion medium), hBO could be established (**Table S1**). Furthermore, we could mature the hBO by culturing them with advanced DMEM/F12 medium containing FGF2, FGF7, FGF10, Y-27632, and A83-01 (differentiation medium) (**Table S1**). Among the growth factors included in the differentiation medium, FGF2 is important for enhancing the expression levels of *ACE2* and *TMPRSS2* (**Fig. S1**). Approximately 100 hBO were present in 50 μL of Matrigel, and the diameter of each hBO was around 100-200 μm (**Fig. 1A**). Transmission electron microscopy (TEM) images showed the presence of cilia and goblet cells (**Fig. 1B, Fig. S2**). The *ACE2* and *TMPRSS2* expression levels in hBO were higher than in cryopreserved adult hBEpC (**Fig. 1C**). Immunohistochemical analysis showed that ACE2 was expressed in part of the outer edge of hBO, while TMPRSS2 was expressed in part of the outer edge and lumen (**Fig. 1D, Fig. S3**). Because bronchi are composed of basal, ciliated, goblet, and club cells, a gene expression analysis of markers specific to these four cell types was performed. The gene expression levels of basal, ciliated, goblet, and club cell markers in hBO were higher than in cryopreserved adult hBEpC (**Figs. 1E-1H**). Consistently, hBO were also positive for α-tubulin, CC10, mucin 5AC, and KRT5 (**Fig. 1I**). The outer edge and lumen of hBO was positive for KRT5 and acetylated α-tubulin, respectively. These results suggest that basal cells are positive for both ACE2 and TMPRSS2, but ciliated cells are positive only for TMPRSS2. Based on these observations, we succeeded in generating expandable and functional hBO from cryopreserved adult hBEpC.

**Figure 1.**
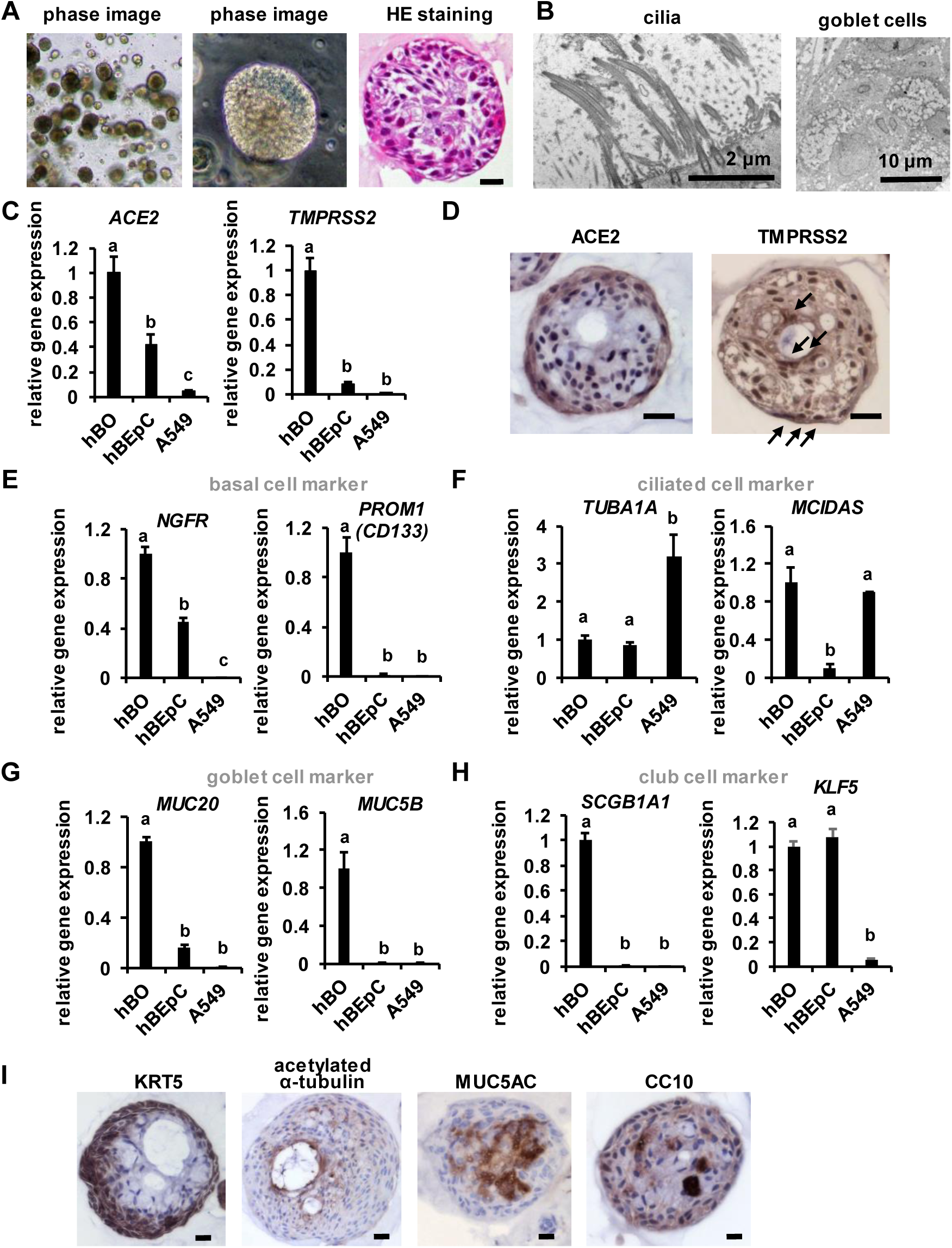
Generation of human bronchial organoids. (A) Phase and HE staining images of human bronchial organoids (hBO). Scale bar = 20 μm. (B) A TEM image of hBO. Larger images are shown in Fig. S2. (C) The gene expression levels of *ACE2* and *TMPRSS2* in hBO and hBEpC were examined by qPCR. The gene expression levels in hBO were normalized to 1.0. (D) The expressions of ACE2 and TMPRSS2 were examined by immunohistochemistry. Scale bar = 20 μm. (E-H) The gene expression levels of basal (E), ciliated (F), goblet (G), and club (H) cell markers in hBO, hBEpC, and A549 cells were examined by qPCR. The gene expression levels in hBO were normalized to 1.0. (I) Immunohistochemistry analysis of KRT5 (basal cell marker), acetylated α-tubulin (ciliated cell marker), Mucin 5AC (goblet cell marker), and CC10 (club cell marker) in hBO. Scale bars = 20 μm. All data are represented as means ± SD (*n* = 3). Statistical significance was evaluated by one-way ANOVA followed by Tukey’s post-hoc tests. Groups that do not share the same letter are significantly different from each other (*p* < 0.05).

### RNA-seq analysis of human bronchial organoids

RNA-seq analysis was performed to further characterize hBO. A heat map, principal component analysis (PCA), and scatter plot of gene expression profiles all show hBO is closer to hBEpC than to A549 cells (**Figs. 2A-2C**). A second heat map of bronchial epithelial markers showed that hBO expressed bronchial epithelial markers more strongly than did hBEpC or A549 cells (**Fig. 2D**). These results suggest that hBO have higher bronchial functions than A549 cells or hBEpC.

**Figure 2.**
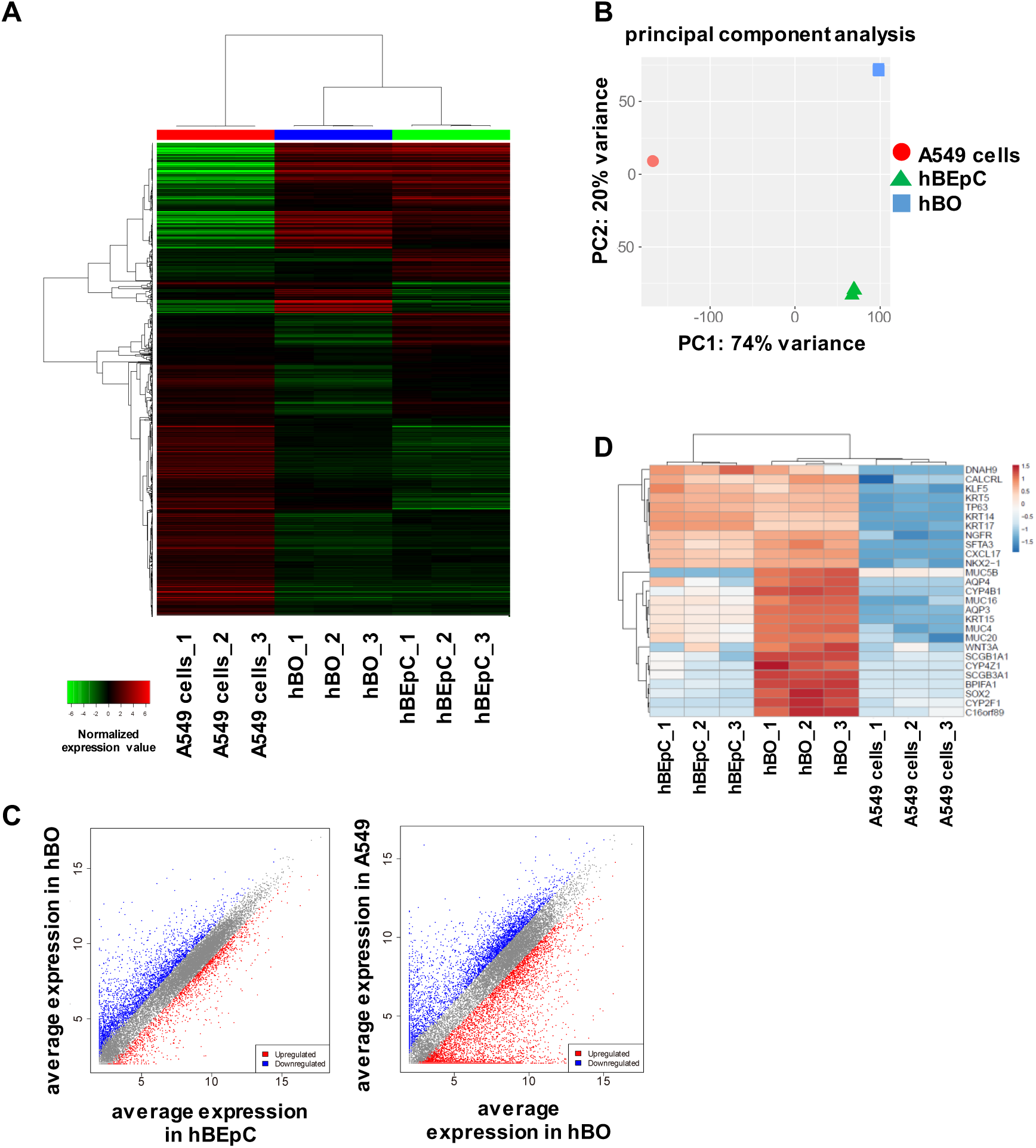
Global gene expression profile of human bronchial organoids. RNA seq analysis was performed in A549 cells (A549), human bronchial organoids (hBO), and human bronchial epithelial cells (hBEpC). (A) Hierarchical clustering analysis of 2,000 variable genes was performed. (B) Principal component analysis (PCA) in “A549”, “hBO”, and “hBEpC”. (C) A scatter plot in “A549”, “hBO”, and “hBEpC”. (D) A clustering analysis of bronchial markers was performed.

### SARS-CoV-2 infection experiments using human bronchial organoids

Next, we investigated whether hBO can be applied to SARS-CoV-2 research. HBO were infected with SARS-CoV-2 and then cultured in differentiation medium for 5 days (**Fig. 3A**). We observed a significant accumulation of LDH in the culture medium of infected hBO (**Fig. 3B**), suggesting that cytotoxicity was caused by the infection. At day 5 after the infection, viral gene expression in infected hBO was clearly detected (**Fig. 3C**). IHC analysis showed that S protein-positive cells were observed in part of the outer edge of hBO (**Fig. 3D**). In addition, S protein co-localized with KRT5 (**Fig. 3E**), but not with CC10 (**Fig. S4**), suggesting that SARS-CoV-2 infected and replicated in basal cells. Overall, these results indicate that SARS-CoV-2 can infect and replicate in hBO.

**Figure 3.**
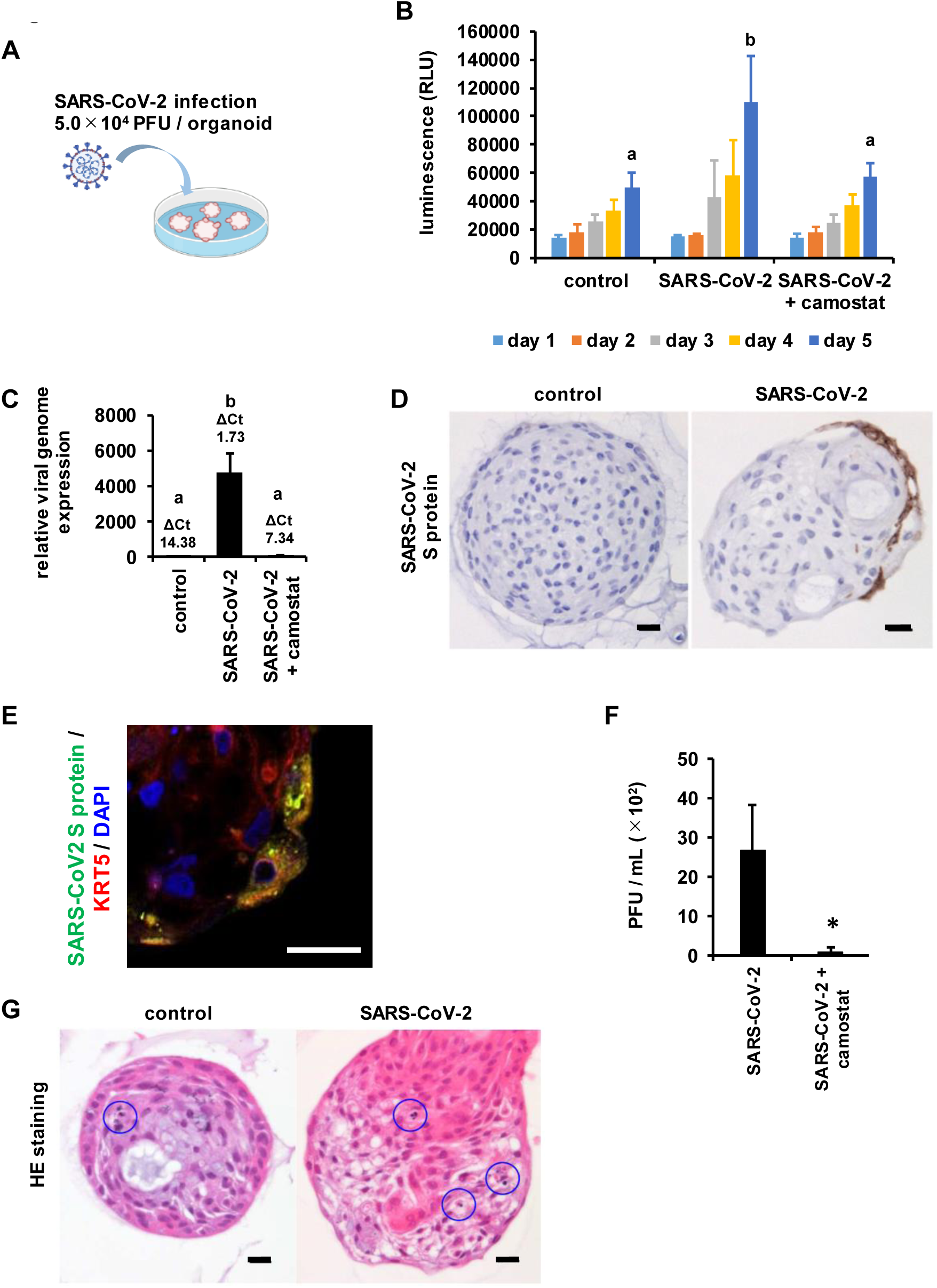
SARS-CoV-2 infection experiments in human bronchial organoids. (A) hBO were infected with SARS-CoV-2 (5.0×10^4^ PFU/well) in the presence or absence of 10 μM camostat and then cultured with differentiation medium for 5 days. (B) At days 1, 2, 3, 4, and 5 after the infection, an LDH assay was performed. (C) The viral genome expression levels in uninfected organoids (control), infected organoids (SARS-CoV-2), and infected organoids treated with camostat (SARS-CoV-2 + camostat) were examined by qPCR. The gene expression levels in control were normalized to 1.0. Statistical significance was evaluated by one-way ANOVA followed by Tukey’s post-hoc tests. Groups that do not share the same letter are significantly different from each other (*p* < 0.05). (D) The expression of SARS-CoV-2 Spike protein was examined by immunohistochemistry. Scale bar = 20 μm. (E) The expression of SARS-CoV-2 Spike protein and KRT5 was confirmed by immunofluorescence staining. Nuclei were counterstained with DAPI. Scale bar = 20 μm. (F) The amount of infectious virus in the supernatant was measured. Statistical analysis was performed using the unpaired two-tailed Student’s *t*-test (* *p* < 0.05). (G) HE staining images of uninfected organoids (control) and infected organoids (SARS-CoV-2) are shown. Blue circles show the existence of pyknotic cells. Scale bar = 20 μm. All data are represented as means ± SD (*n* = 3).

Currently, clinical trials using camostat, favipiravir, nafamostat, chloroquine, ritonavir/lopinavir, and remdesivir are underway around the world to develop therapeutic agents for COVID-19. However, the evaluation of these drugs using *in vitro* lung models is rare. Therefore, in this study, the effect of camostat, an inhibitor of TMPRSS2, was examined using our hBO because camostat was demonstrated as a promising candidate in cell culture models ^3^. Upon camostat treatment, the amount of SARS-CoV-2 viral genome was reduced to 2% of untreated infected hBO (**Fig. 3C**). In addition, LDH release from infected hBO was significantly reduced after camostat treatment (**Fig. 3B**). Finally, we examined the culture supernatants of infected hBO. We found that infectious virus was significantly observed in infected hBO, but the production of infectious virus was impaired by treatment with camostat (**Fig. 3F**). Collectively, our data indicated that hBO can secrete infectious virus into the culture medium, suggesting that our hBO system can investigate the entire life cycle of SARS-CoV-2.

Next, we examined the pathological effects of the SARS-CoV-2 infection. We observed that the number of pyknotic cells seemed to increase with the infection (**Fig. 3G**). In addition, the expression levels of type I IFN (IFN-I) and IFN-stimulated genes were moderately increased after SARS-CoV-2 infection (**Fig. S5A**). Furthermore, SARS-CoV-2 infection did not change the gene expression levels of *ACE2* or *TMPRSS2* (**Fig. S5B**). From the above, these results suggest that hBO can be used to reproduce SARS-CoV-2-induced pulmonary disorder and to evaluate the effect of therapeutic agents.

### RNA-seq analysis of SARS-CoV-2-infected human bronchial organoids

RNA-seq analysis was performed to investigate the effects of SARS-CoV-2 infection and camostat treatment in detail. A heat map shows that the gene expression profile of SARS-CoV-2-infected hBO is closer to SASR-CoV-2-infected hBO treated with camostat than uninfected hBO (**Fig. 4A**). A PCA and scatter plot of the gene expressions agree with this finding (**Figs. 4B, C**). Additionally, SARS-CoV-2 infection increased the expression levels of IFN-I signaling-related genes in hBO (**Fig. 4D**). PGSEA applied on GO biological process gene sets shows that the expression levels of genes involved in positive regulation of immune effector process, regulation of inflammatory response, interferon-gamma production, and positive regulation of acute inflammatory response were increased by SARS-CoV-2 infection and that this increase was suppressed by camostat treatment (**Fig. 4E**). These results indicated that SARS-CoV-2 infection induces IFN-I signaling-related genes and that camostat treatment reversed this phenotype.

**Figure 4.**
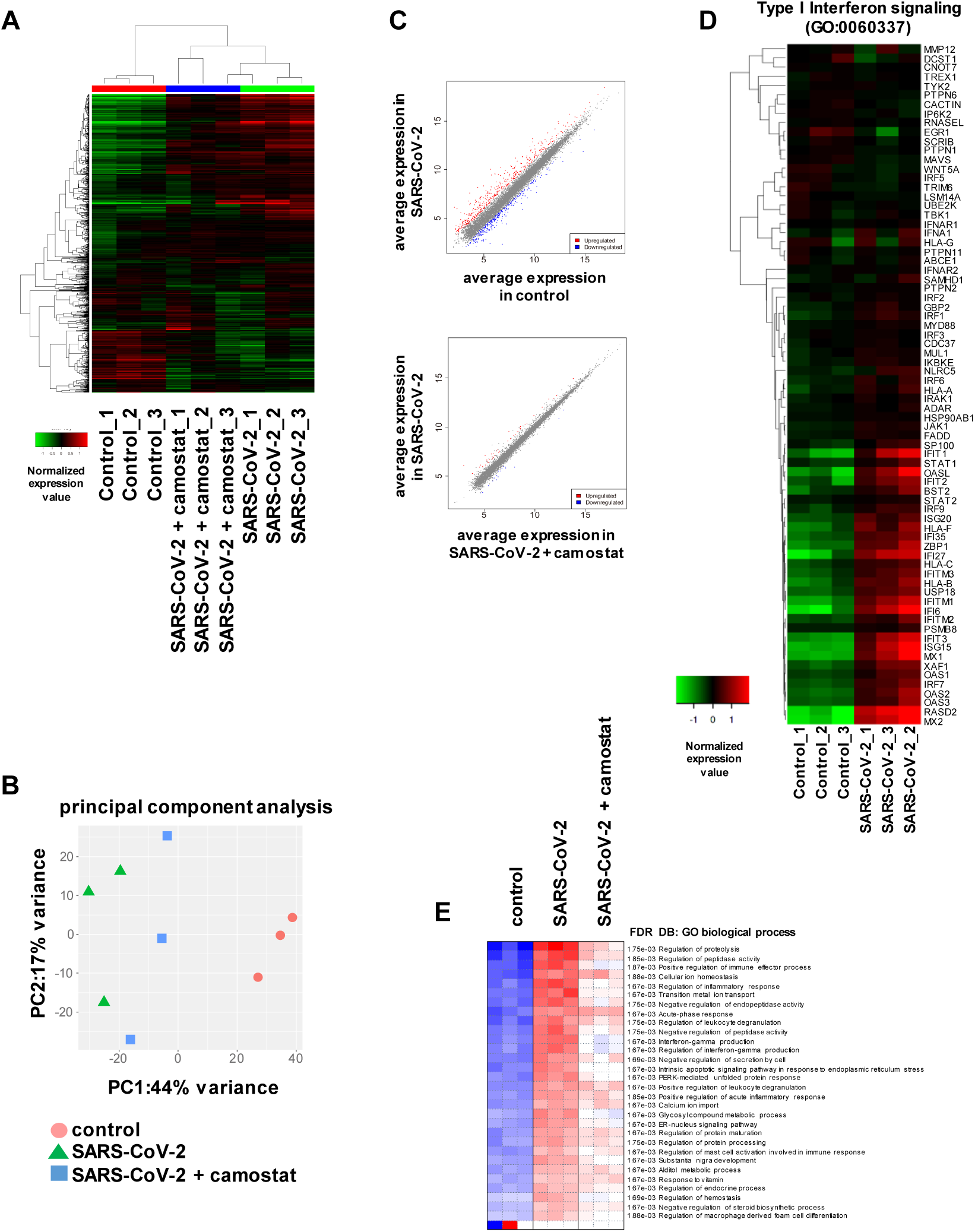
Global gene expression profile of infected human bronchial organoids. RNA seq analysis of uninfected hBO (control), SARS-CoV-2-infected hBO (SARS-CoV-2), and SARS-CoV-2-infected hBO treated with camostat (SARS-CoV-2 + camostat). (A) A clustering analysis of 2,000 variable genes was performed. (B) Principal component analysis (PCA) in “control”, “SARS-CoV-2”, “SARS-CoV2 + camostat”. (C) A scatter plot in “control”, “SARS-CoV-2”, “SARS-CoV2 + camostat”. (D) A heat map of IFN-I signaling-related genes in “control” and “SARS-CoV-2” is shown. (E) PGSEA (Parametric Gene Set Enrichment Analysis) applied on GO biological process gene sets was performed.

## Discussion

In this study, we succeeded in generating hBO from cryopreserved adult hBEpC and applied it to SARS-CoV-2 research. We confirmed that SARS-CoV-2 could infect and replicate in these cells and that camostat could suppress the replication. If small airway organoids and alveolar organoids can be produced from cryopreserved adult human small airway epithelial cells and alveolar epithelial cells, respectively, the infection and replication of SARS-CoV-2 in each part of the lung can be evaluated. Recently, it was shown that SARS-CoV-2 infection and replication can be observed in kidney ^12^, liver ductal ^13^, and gut organoids ^14^. By comparing the infection and replication ability of SARS-CoV-2 in these organoids, the sensitivity of SARS-CoV-2 in each organ could be compared.

The incorporation of mechanical stress into our organoid system could improve the accuracy of SARS-CoV-2 research. The human airway is always exposed to shear stress due to air flow. It has been reported that a functional *in vitro* lung model can be generated using a device capable of medium perfusion and expansion/contraction (organ-on-a-chip) ^15^. Recently, it was reported that the infection and replication of SARS-CoV-2 can be observed by culturing primary human lung airway epithelial basal stem cells and pulmonary microvascular endothelial cells on a chip device ^16^. By applying our hBO to a similar device, we may be able to construct an *in vitro* bronchi model that more closely mimics the living body.

Nevertheless, even in its current condition, we used our hBO to evaluate the efficacy of a COVID-19 therapeutic agent and observe the cytotoxicity and innate immune responses caused by the SARS-CoV-2. Moreover, we could clarify the localization of SARS-CoV-2 in the hBO. These results could be obtained because hBO reproduce the cell populations and functions of the bronchi. Our data of RNA-seq analysis would give us a change to understand the virus affects cellular and bronchial functions.

Finally, we showed that camostat has a positive effect against COVID-19 infection in hBO, demonstrating its usefulness for COVID-19 drug discovery. Similar studies on experimental COVID-19 drugs including those currently undergoing clinical trials should be considered. Furthermore, because hBO can be generated from commercially available cryopreserved hBEpC quickly (10 days) and at large scale, we expect hBO will shorten the search for effective COVID-19 agents.

## Materials and Methods

### Human bronchial organoid culture

Normal human bronchial epithelial cells (hBEpC, Lonza) were suspended in 10 mg/ml cold Matrigel growth factor reduced (GFR) basement membrane matrix. 50 μL drops of cell suspension were solidified on pre-warmed Nunc cell-culture treated multidishes (24-well plate) at 37°C for 10 min, and then 500 μL of expansion medium (composition is shown in **Table S1**) was added to each well. The expansion medium was changed every 2 days. HBO were passaged every 10-12 days. For passaging, the hBO were suspended in 1 mL of 0.5 mM EDTA/PBS (Nacalai tesque) and mechanically sheared using a P1000 pipette tip. Then, 2 mL TrypLE Select (Thermo Fisher Scientific) was added to the suspension. After incubating for 5 min at room temperature, the hBO were again mechanically sheared using a P1000 pipette tip. 7 mL of expansion medium was added, and the organoid suspension tubes were centrifuged at 400 rpm. Organoid fragments were re-suspended in cold expansion medium and seeded as above. HBO were passaged every 10 days. To mature the hBO, the expanded hBO were cultured with differentiation medium (composition is shown in **Table S1**) for 5 days. HBO can be cryopreserved by using STEM-CELLBANKER GMP grade (TaKaRa Bio).

### A549 culture

A549 cells were cultured with Ham’s F12 medium (Thermo Fisher Scientific) containing 10% FBS, 1×GlutaMAX (Thermo Fisher Scientific), and penicillin-streptomycin. A549 cells were passaged every 4 days.

### SARS-CoV-2 preparation

The SARS-CoV-2 strain used in this study (SARS-CoV-2/Hu/DP/Kng/19-020) was obtained from the Kanagawa Prefectural Institute of Public Health. SARS-CoV-2 was isolated from a COVID-19 patient in Japan (GenBank: LC528232.1). The isolation and analysis of the virus will be described elsewhere (manuscript in preparation). The virus was plaque-purified and propagated in Vero E6 cells. SARS-CoV-2 was stored at −80°C. All experiments including virus infections were done in the biosafety level facility at Osaka University strictly following regulations.

### SARS-CoV-2 infection and drug treatment

Approximately 100 organoids were infected with 5.0×10^4^ PFU of SARS-CoV-2 in a 24-well plate containing 500 uL differentiation medium. One-half of the differentiation medium containing SARS-CoV-2 was replaced with fresh differentiation medium every day. At 5 days after the infection, the hBO and their supernatant were collected. In the drug treatment experiments, the infected hBO were cultured with differentiation medium containing 10 μM camostat (Sigma-Aldrich) for 5 days.

### SARS-CoV-2 virus plaque assays

VeroE6/TMPRSS2 cells (JCRB1819, JCRB Cell Bank) ^17^ were seeded on 12 well plates (1.7×10^5^ cells/well) and incubated for 24 hr. The culture supernatants serially diluted by medium were inoculated and incubated for 2 hr. Culture medium was removed, fresh medium containing 1% methylcellulose (1.5mL) was added, and the culture was further incubated for 3 days. The cells were fixed with 4% Paraformaldehyde Phosphate Buffer Solution (Nacalai Tesque) and plaques were visualized by using a Crystal violet.

### Quantitative PCR

Total RNA was isolated from hBO using ISOGENE II (NIPPON GENE). cDNA was synthesized using 500 ng of total RNA with a Superscript VILO cDNA synthesis kit (Thermo Fisher Scientific). Real-time RT-PCR was performed with the SYBR Green PCR Master Mix (Applied Biosystems) using a StepOnePlus real-time PCR system (Applied Biosystems). The relative quantitation of target mRNA levels was performed by using the 2-ΔΔCT method. The values were normalized by those of the housekeeping gene, *glyceraldehyde 3-phosphate dehydrogenase* (*GAPDH*). The PCR primer sequences are shown in **Table S2**.

The SARS-CoV-2 primer and probe sets were obtained from Integrated DNA Technologies (IDT, 10006606).

### Ultrathin section transmission electron microscopy (TEM)

HBO fixed in phosphate buffered 2% glutaraldehyde, and subsequently post-fixed in 2% osmium tetra-oxide for 2 hr at 4°C. After fixation, they were dehydrated in a graded series of ethanol and embedded in the epoxy resin. Ultrathin sections were cut and then stained with uranyl acetate and lead staining solution and were examined using an electron microscope (HITACHI H-7600) at 100 kV.

### Histopathology and immunofluorescence

Fixed bronchial organoid samples were processed and embedded in paraffin. Then they were cut into 2 μm-thick sections. The sections were deparaffinized, rehydrated, and stained with hematoxylin and eosin (HE). The sections were then examined using a microscope (BX53 microscope with DP73 camera, Olympus Corporation).

For the immunohistochemical stain assay, the formalin-fixed and paraffin-embedded bronchial organoid samples were treated with pH 6.0 citrate buffer for 30 sec at 125°C in a pressure cooker (Dako Japan) as antigen retrieval. Sections were incubated with each antibody (**Table S3**), followed by Histofine Simple Stain MAX-PO (Nichirei Biosciences). The sections were visualized using Peroxidase Stain DAB Kit (Nacalai Tesque) before counterstaining with Meyer’s hematoxylin.

For the double immunofluorescence staining assay, the sections were deparaffinized and subjected to antigen retrieval by treating them with 0.5% trypsin for 30 min. Then the sections were blocked by 5% skim milk with albumin obtained from Bovine Serum Cohn Fraction V, pH 7.0 (Wako Pure Chemical Industries), in PBS for 30 min at room temperature to avoid non-specific reactions. The sections were then incubated with primary antibody (**Table S3**) overnight at 4°C, washed, and incubated with secondary antibody (**Table S3**) for 1 h at room temperature. After washing with PBS, the specimens were mounted with glycerol. All observations were performed using the BX53 fluorescence microscope with a DP73 camera equipped with a suitable filter set (red filter with excitation range 530-550 nm and emission range 575 nm, and green filter with excitation range 470-495 nm and emission range of 510 nm).

### RNA-seq

Total RNA was prepared using the RNeasy Mini Kit (Qiagen). RNA integrity was assessed with a 2100 Bioanalyzer (Agilent Technologies). Library preparation was performed using a NEBNext Ultra II Directional RNA Library Prep Kit for Illumina (NEB) or a TruSeq stranded mRNA sample prep kit (Illumina) according to the manufacturer’s instructions. Sequencing was performed on an Illumina NextSeq500 or NovaSeq6000 platform in 152- or 101-base single-end mode, respectively. Fastq files were generated using bcl2fastq2. Adapter sequences were trimmed from the raw reads by cutadapt ver 2.7. The trimmed reads were mapped to the human reference genome sequences (hg19) using HISAT2 ver 2.1.0. The raw counts were calculated using featureCounts ver 2.0.0 and used for heatmap visualization with integrated differential expression and pathway analysis (iDEP, (http://ge-lab.org/idep/)) ^18^. Access to raw data concerning this study was submitted under Gene Expression Omnibus (GEO) accession number GSE150819.

### LDH assay

After the SARS-CoV-2 infection, the release of LDH was monitored from an aliquot of 250 μL supernatant using the LDH-Glo cytotoxicity assay (Promega) according to the manufacturer’s instructions. The absorbance was determined with a Bio-Rad microplate reader (Bio-Rad, US) at wavelength 490 nm. The release of LDH in uninfected cells was used as a control.

### Statistical analyses

Statistical analysis was performed using the unpaired two-tailed Student’s *t*-test. Statistical significance was evaluated by one-way analysis of variance (ANOVA) followed by Tukey’s or Dunnett’s post hoc tests to compare all groups.

## Abbreviations

2D: two-dimensional
ACE2: angiotensin-converting enzyme 2
CC10: club cell protein 10
FGF: fibroblast growth factor
hBEpC: human bronchial epithelial cells
hBO: human bronchial organoids
IFN-I: type I interferon
IHC: immunohistochemistry
KRT5: keratin 5
LDH: lactate dehydrogenase
PSC: pluripotent stem cell
RdRp: RNA-dependent RNA polymerase
RNA: seq RNA sequencing
SARS-CoV-2: severe acute respiratory syndrome coronavirus 2
TMPRSS2: transmembrane serine proteinase 2
WHO: World Health Organization

## Acknowledgements

The SARS-CoV-2 strain used in this study (SARS-CoV-2/Hu/DP/Kng/19-020) was obtained from Kanagawa Prefectural Institute of Public Health. This research was supported by the iPS Cell Research Fund. The figure 3A was created using Biorender (https://biorender.com). We thank Dr. Tomohiko Takasaki and Dr. Jun-Ichi Sakuragi (Kanagawa Prefectural Institute of Public Health) for providing SARS-CoV-2 strain (SARS-CoV-2/Hu/DP/Kng/19-020), Dr. Misaki Ouchida (Kyoto University) for creating graphical abstract, Dr. Peter Karagiannis (Kyoto University) for critical reading of the manuscript, Dr. Nobihiro Morone (University of Cambridge) for critical discussions, Ms. Sayaka Deguchi (Osaka University) for technical assistance with the qPCR analysis, and Ms. Kazusa Okita and Ms. Eri Kawaguchi for technical assistance with the RNA-seq experiments.

## Author Contributions

TS performed the SARS-CoV-2 experiments and analyses

YI performed the SARS-CoV-2 experiments and analyses

YS performed the immunohistochemical analysis

AS prepared the materials for the SARS-CoV-2 experiments and analyses

DO performed the analysis of RNA-seq data of the infected bronchial organoids

DM collected RNA-seq data of the infected bronchial organoids

SM performed the SARS-CoV-2 experiments and analyses

TK performed the SARS-CoV-2 experiments and analyses

TY collected the RNA-seq data of the bronchial organoids

TO designed the research and performed the SARS-CoV-2 experiments and analyses

KT designed the research, generated the bronchial organoids, performed statistical analysis, and wrote the paper

## Declaration of interests

The authors declare no competing financial interests.

## Supplemental figures

**Figure S1.**
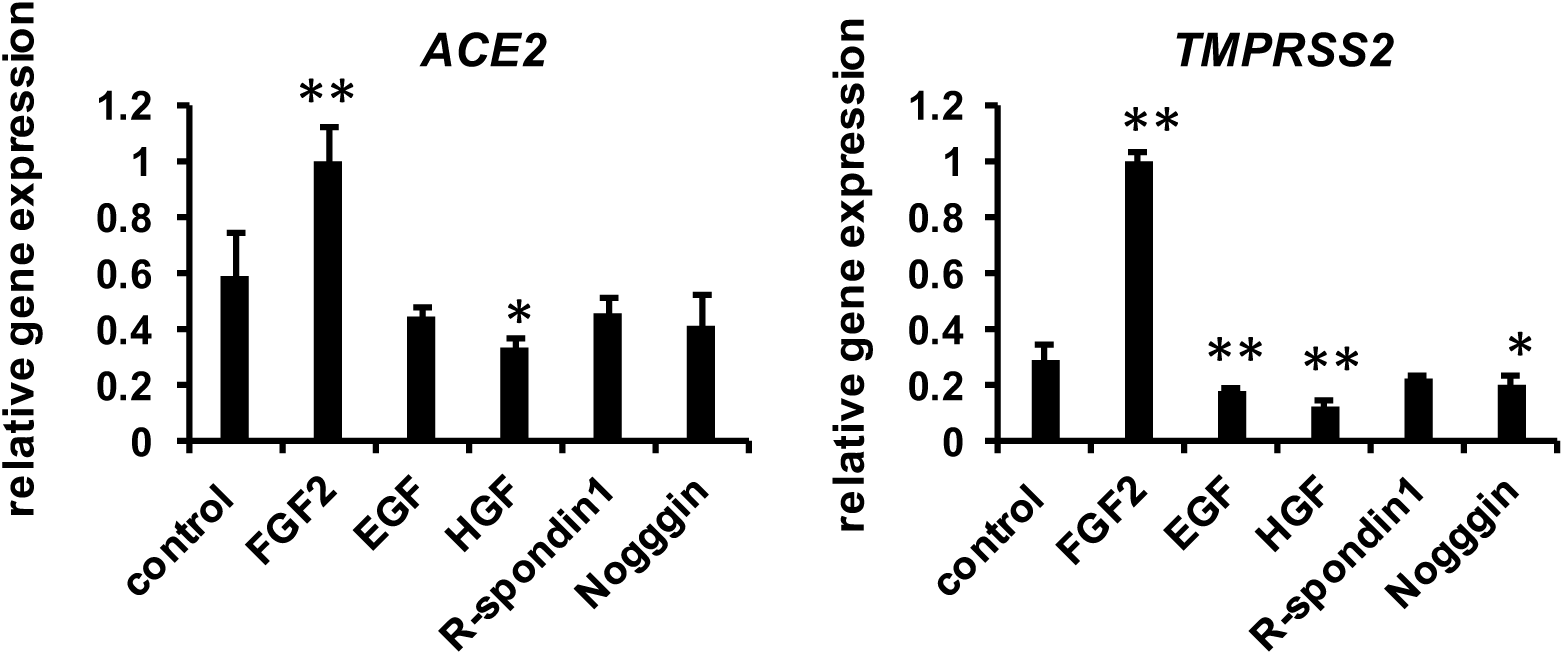
FGF2 promotes the maturation of human bronchial organoids. Expanded hBO were cultured with differentiation medium containing FGF2, EGF, HGF, R-spondin 1, or Noggin for 5 days. The gene expression levels of *ACE2* and *TMPRSS2* were examined by qPCR. The gene expression levels in “FGF2” were normalized to 1.0. All data are represented as means ± SD (*n* = 3). Statistical significance was evaluated by one-way ANOVA followed by Dunnett’s post-hoc tests (* *p* < 0.05, ** *p* < 0.01, as compared with control).

**Figure S2.**
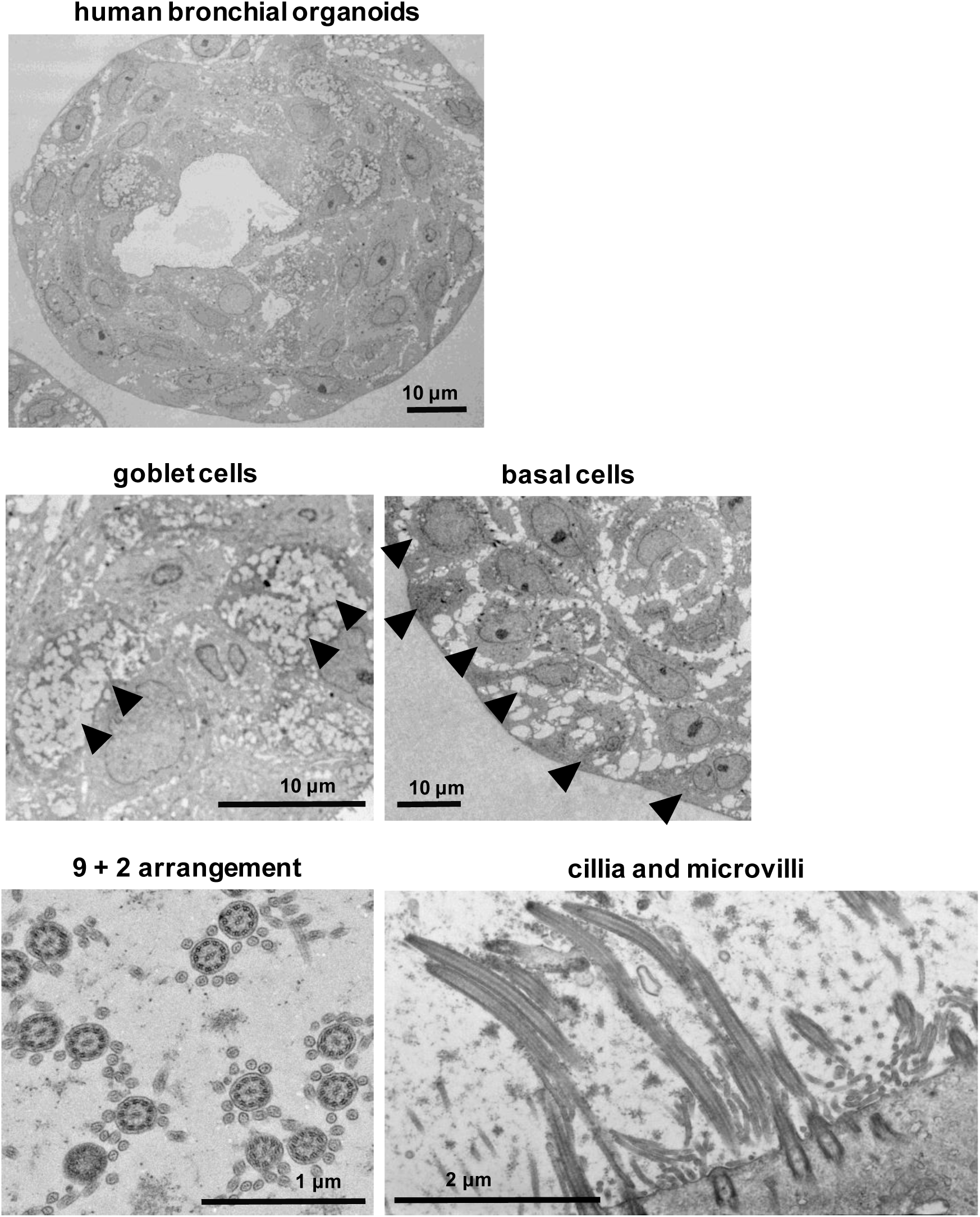
TEM image of human bronchial organoids. The larger TEM images of Fig. 1B are shown. Goblet cells, basal cells, 9+2 arrangement, cilia, and microvilli can be observed in hBO.

**Figure S3.**
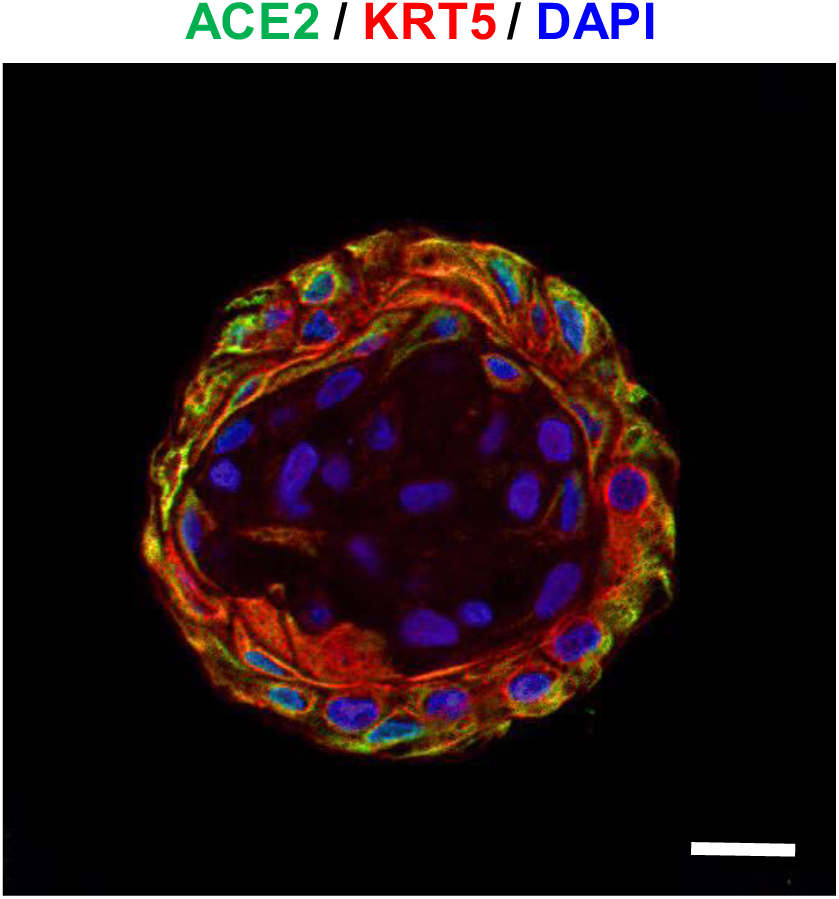
ACE2 is expressed in basal cells. The expression of ACE2 and KRT5 (basal cell marker) in hBO was confirmed by immunofluorescence staining. Nuclei were counterstained with DAPI. Scale bar = 20 μm.

**Figure S4.**
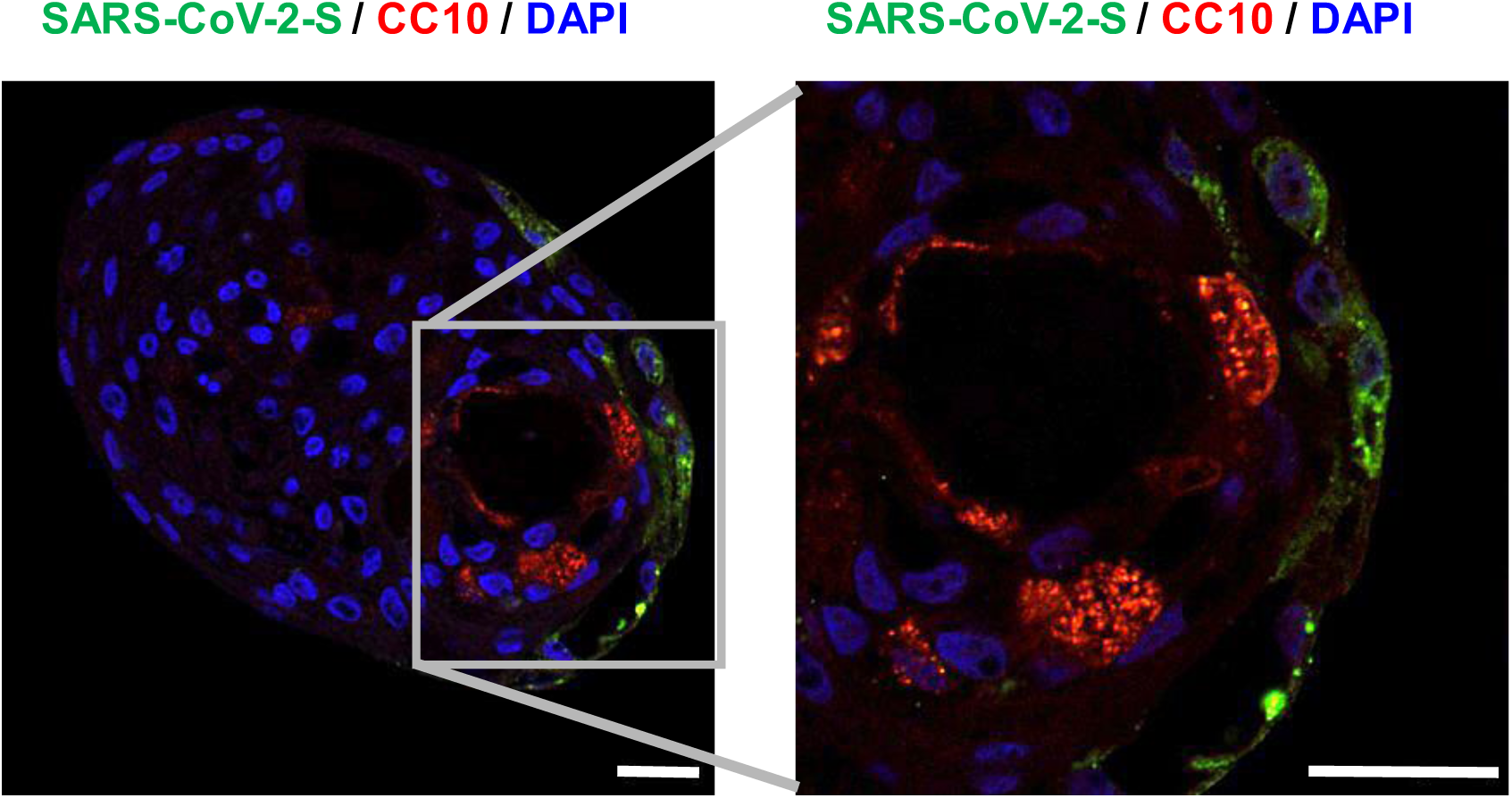
SARS-CoV-2 Spike protein is not expressed in club cells. hBO were infected with SARS-CoV-2 (5.0×10^4^ PFU/well) and then cultured with differentiation medium for 5 days. The expression of SARS-CoV-2 Spike protein and CC10 was confirmed by immunofluorescence staining. Nuclei were counterstained with DAPI. Scale bar = 20 μm.

**Figure S5.**
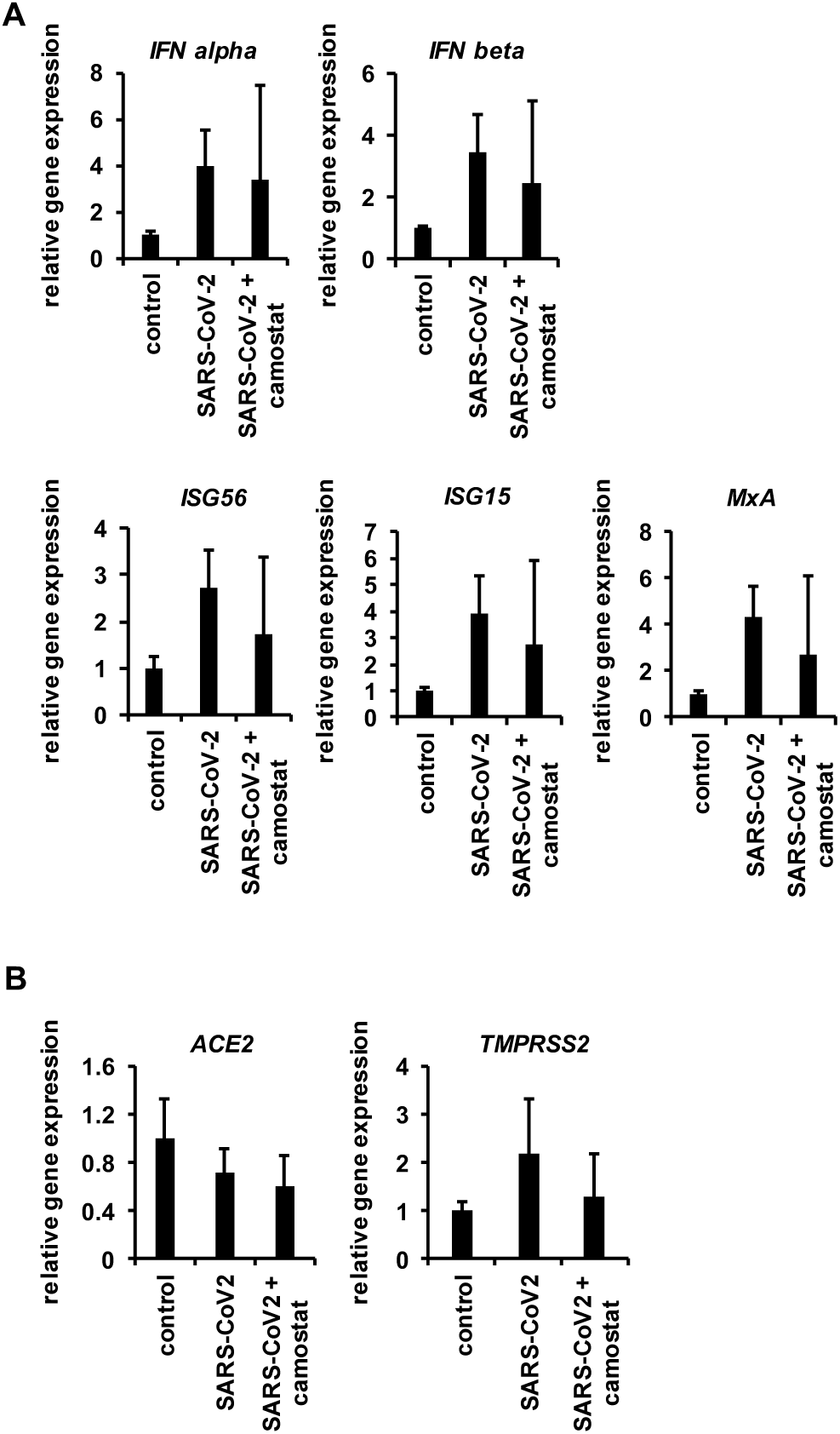
SARS-CoV-2-induced innate immune responses in human bronchial organoids. hBO were infected with SARS-CoV-2 (5.0×10^4^ PFU/well) in the presence or absence of 10 μM camostat and then cultured with differentiation medium for 5 days. (A, B) The gene expression levels of type I interferon (*IFN alpha* and *IFN beta*), interferon-stimulated genes (*ISG56, ISG15*, and *MxA*) (A), and SARS-CoV-2-associated genes (*ACE2* and *TMPRSS2*) (B) in uninfected organoids (control), infected organoids (SARS-CoV-2), and infected organoids treated with camostat (SARS-CoV-2 + camostat) were examined by qPCR. The gene expression levels in control were normalized to 1.0. All data are represented as means ± SD (*n* = 3).

## Tables

**Table S1.**
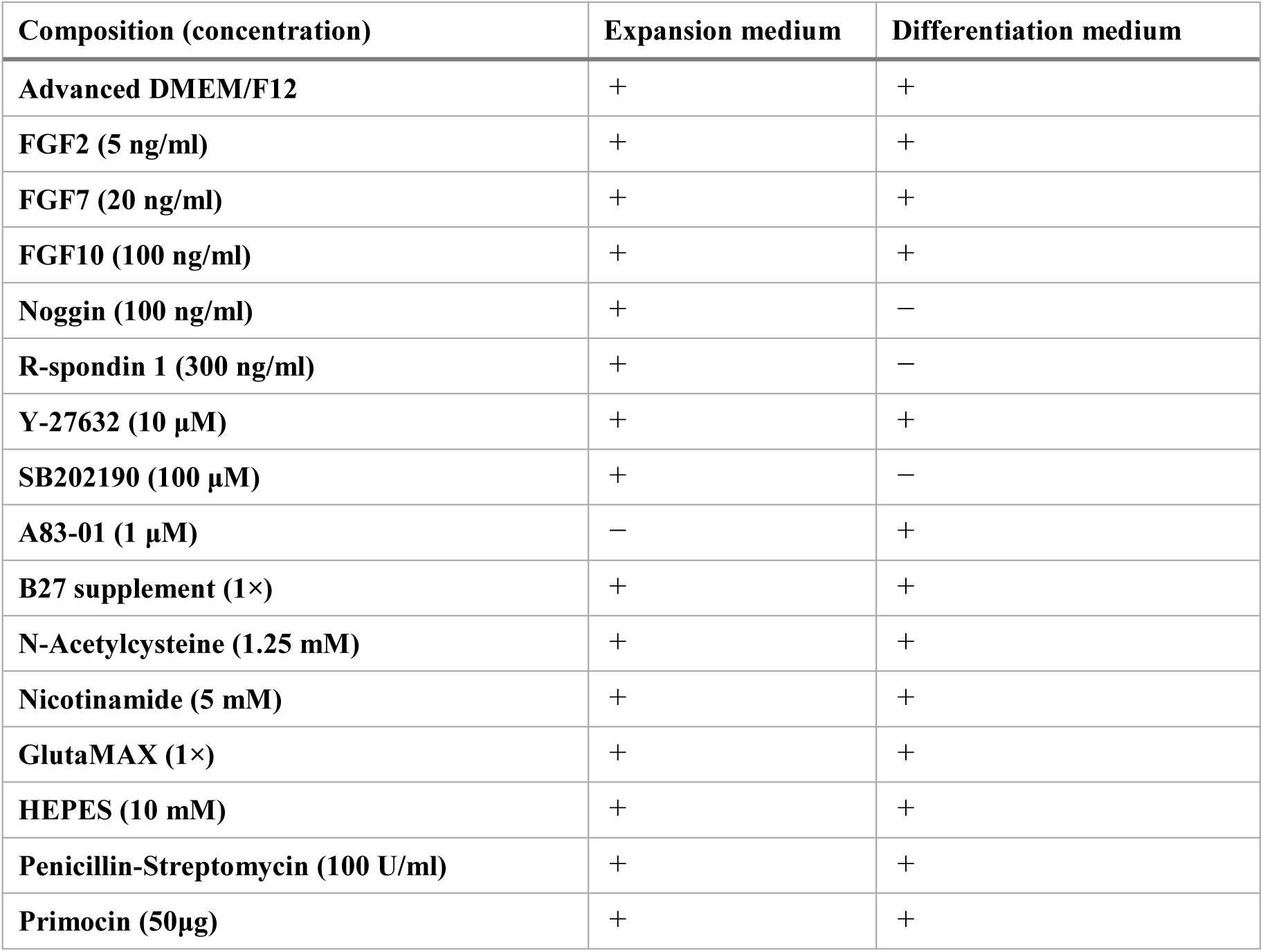
Composition of expansion and differentiation media for human bronchial organoids.

| Composition (concentration) | Expansion medium | Differentiation medium |
| --- | --- | --- |
| Advanced DMEM/F12 | + | + |
| FGF2 (5 ng/ml) | + | + |
| FGF7 (20 ng/ml) | + | + |
| FGF10 (100 ng/ml) | + | + |
| Noggin (100 ng/ml) | + | — |
| R-spondin 1 (300 ng/ml) | + | — |
| Y-27632 (10 μM) | + | + |
| SB202190 (100 μM) | + | — |
| A83-01 (1 μM) | — | + |
| B27 supplement (1×) | + | + |
| N-Acetylcysteine (1.25 mM) | + | + |
| Nicotinamide (5 mM) | + | + |
| GlutaMAX (1×) | + | + |
| HEPES (10 mM) | + | + |
| Penicillin-Streptomycin (100 U/ml) | + | + |
| Primocin (50μg) | + | + |

**Table S2.**
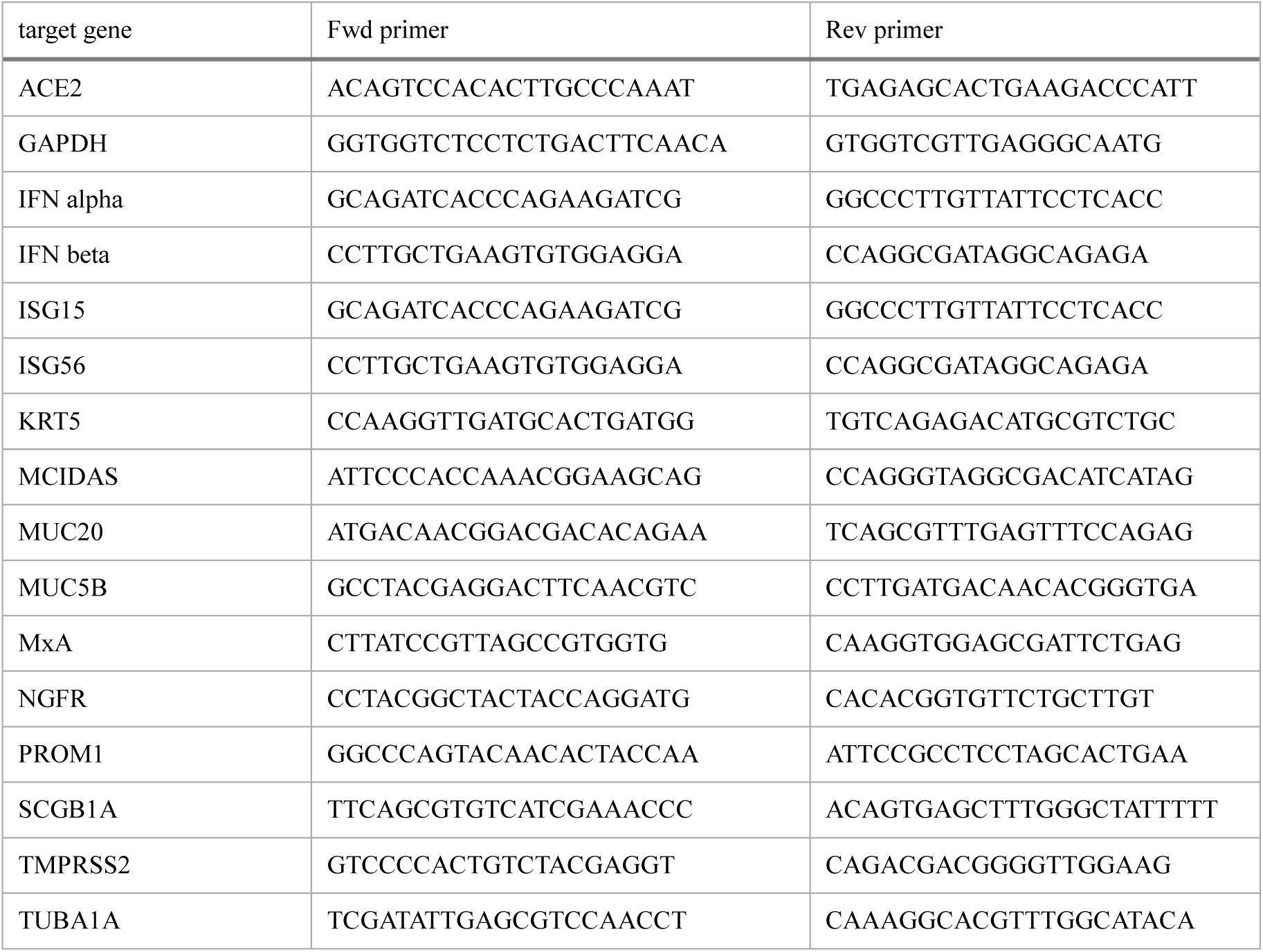
Primer list.

**Table S3.** Antibody list.

| Antigen (clone number) | catalog number | company |
| --- | --- | --- |
| ACE2 | PGI-21115-1-AP-150 | Proteintech |
| acetylated $\alpha$ tubulin (6-11B-1) | sc-23950 | Santa Cruz Biotechnology |
| CC10 (E-11) | sc-365992 | Santa Cruz Biotechnology |
| cytokeratin 5 (RCK103) | sc-32721 | Santa Cruz Biotechnology |
| mucin 5AC (45M1) | sc-21701 | Santa Cruz Biotechnology |
| SARS-CoV/SARS-CoV-2 (COVID-19)<br>spike protein (1A9) | GTX632604 | GeneTex |
| TMPRSS2 (H-4) | sc-515727 | Santa Cruz Biotechnology |

## Notes

https://www.ncbi.nlm.nih.gov/geo/query/acc.cgi?acc=GSE150819

### Competing Interest Statement

The authors have declared no competing interest.

